# Macrophage ferroptosis inhibits *Aspergillus* conidial killing in lung transplantation

**DOI:** 10.1101/2025.03.13.643092

**Authors:** Efthymia Iliana Matthaiou, Amer Ali Abd El-Hafeez, Husham Sharifi, Paulami Chatterjee, Matthew Zinter, Patrik Johansson, Ekroop Dhillon, Wayland Chiu, Jin Qian, Brian Shaller, Jiwoon Chang, Shravani Pasupneti, Carlos Hernandez Borges, Sarah Omar, Annika Enejder, Gundeep Dhillon, Brice Gaudilliere, Jarrod Fortwendel, Jatin M. Vyas, Joe L. Hsu

## Abstract

Immune suppression heightens the risk for fungal infections, but the mechanisms that result in clinical disease are poorly understood. Here we demonstrate that macrophage ferroptosis, an iron-dependent form of regulated cell death, inhibits *Aspergillus fumigatus* (*Af*) killing. In a mouse tracheal transplant model of *Af* infection, we observed an increase in macrophage lipid peroxidation, a decreased expression of negative ferroptosis regulators *Gpx4* and *Slc7a11*, and an increase in positive regulators *Ptgs2* and *Nox2*, relative to syntransplants. Depletion of macrophages in transplant recipients decreased *Af* invasion. *In vitro*, iron overload reduced macrophage viability and decreased their capability to kill *Af* spores, through a decrease in lysosomal acidification and lysosomal loss. Treatment with ferrostatin-1, a ferroptosis inhibitor, and deferasirox (an iron chelator) restored *Af* killing. Ferroptotic alveolar macrophages isolated from lung transplant patients also showed a decreased ability to kill *Af* spores and the patients’ bronchoalveolar lavage was characterized by higher iron levels and markers of ferroptotic stress compared to non-lung transplants. These characteristics were strongly correlated with a clinical history of fungal infections, independent of immune suppressive medications. Our findings indicate that macrophage ferroptosis augments the risk of invasive aspergillosis, representing a novel mechanism for host immune dysfunction.

**Graphical Abstract:** Schematic of proposed mechanism underlying ferroptosis induced immune dysregulation and increased *Af* invasion in lung transplantation.

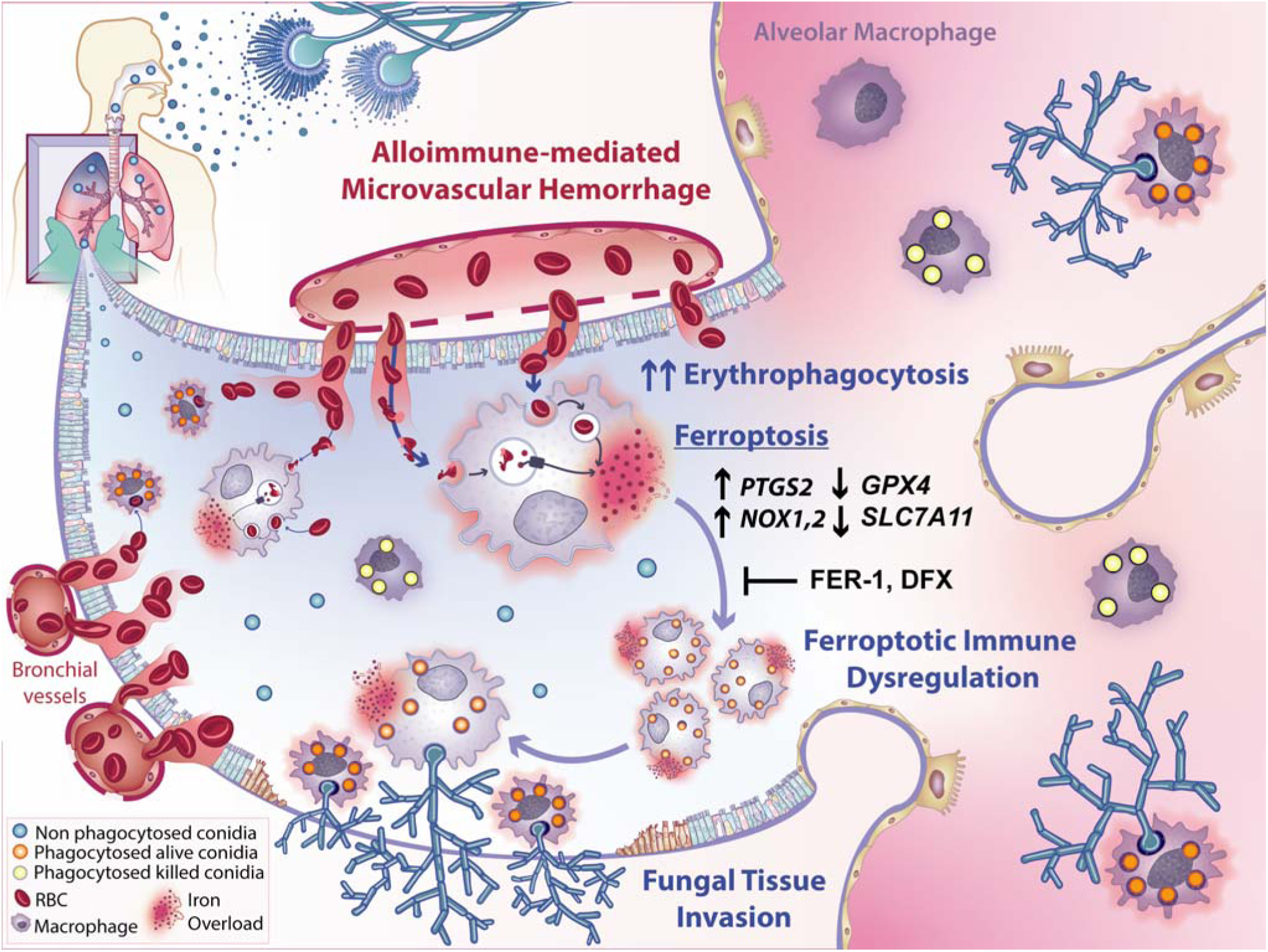

## Introduction

Lung transplantation is a life-saving treatment for persons with several end-stage pulmonary diseases. However, survival may be diminished by infections with *Aspergillus fumigatus (Af),* a ubiquitous mold that releases airborne spores (conidia), and is estimated worldwide to cause 250,000 cases of invasive aspergillosis annually.(1–4) The emergence of *Af* multidrug resistance complicates treatment and leads to higher mortality, which highlights the need for the development of new therapeutic strategies.(5) Among lung transplant recipients (LTRs), up to one in three suffers from an *Af-*related pulmonary disease.(6, 7) This can range from invasive infections of the transplanted airway anastomosis and invasive pulmonary aspergillosis to airway colonization, which in turn accelerates chronic lung allograft dysfunction, the most common cause of death after lung transplant.(7–10) While immunologically vulnerable populations are known to be at risk for fungal infections, the underlying mechanisms that result in a wide array of clinical outcomes are greatly understudied. In persons with hematologic malignancies and after hematopoietic cell transplant, neutropenia is a well-described risk factor for *Af* infections, but in LTRs, severe neutropenia is less commonly seen.(11) Thus, other factors that lead to the increased risk of *Af* infections need to be elucidated and significant improvement is needed in our understanding of the underlying cellular and molecular immunologic responses that lead to effective clearance, colonization, or capitulation to *Af* in LTRs.

To better understand the transplanted host-*Af* relationship, we developed a murine orthotopic tracheal transplant (OTT) model of *Af* infection.(12–14) This model mimics the invasive airway anastomotic infections that occur post lung transplant.(14) We have previously shown that alloimmune-mediated microhemorrhage increases graft iron, which enhances *Af* virulence.(14) However, the interactions among immunity, iron overload and *Af* infection are still poorly understood. Alveolar macrophages (mφs) are the first line of defense against *Af* conidia.(15) They recognize and phagocytose conidia, which are transferred into the phagolysosome for enzymatic killing that requires an acidic pH.(16) However, their role in containing *Af* invasion remains unclear.(15, 17) Mφs are also central in maintaining iron homeostasis and display an iron-dependent increase in recruitment in the OTT model.(14) While mφs are capable of recycling iron, they are susceptible to ferroptosis during increased erythrophagocytosis.(18)

Ferroptosis is a form of regulated cell death (RCD) that results from the production of iron-catalyzed toxic reactive oxygen species (ROS).(18–23) Ferroptosis is characterized by mitochondrial contraction and lysosomal leakage, but differs from other forms of RCD, in that it is independent of caspases, and does not involve chromatin aggregation or disruption of the cell membrane and nuclei remain intact.(18–23) Five defining characteristics of ferroptosis are: (i) iron overload; (ii) the accumulation of ROS; (iii) the presence of lipid peroxidation; (iv) a central role for glutathione peroxidase 4 (GPX4) and (v) inhibition by ferrostatin-1 (FER-1).(18, 23) GPX4 is a protective anti-ferroptotic enzyme that detoxifies lipid peroxides at the expense of reduced glutathione.(22, 23) Ferroptosis, first recognized in cancer, is now known to contribute to Alzheimer’s and Parkinson’s diseases, ischemia reperfusion injury, atherosclerosis, acute kidney injury and response to acute hemorrhage.(20, 24) Given the progressive microhemorrhage and erythrophagocytosis observed in the OTT model, we posited that lung transplant mφs undergo ferroptotic stress, which induces tissue damage, an inability to clear *Af* infection, and fungal invasion. The following studies were designed to test this interpretation and define the underlying mechanisms. We first evaluated the role of ferroptosis in *Af* invasion using the OTT model. Next, we defined the impact of ferroptotic stress on mφ conidial killing *in vitro*. Finally, we validated our preclinical results in bronchoalveolar lavage (BAL) samples collected from lung transplant and non-lung transplant patients (see study schematic Figure 1A).

**Figure 1.**
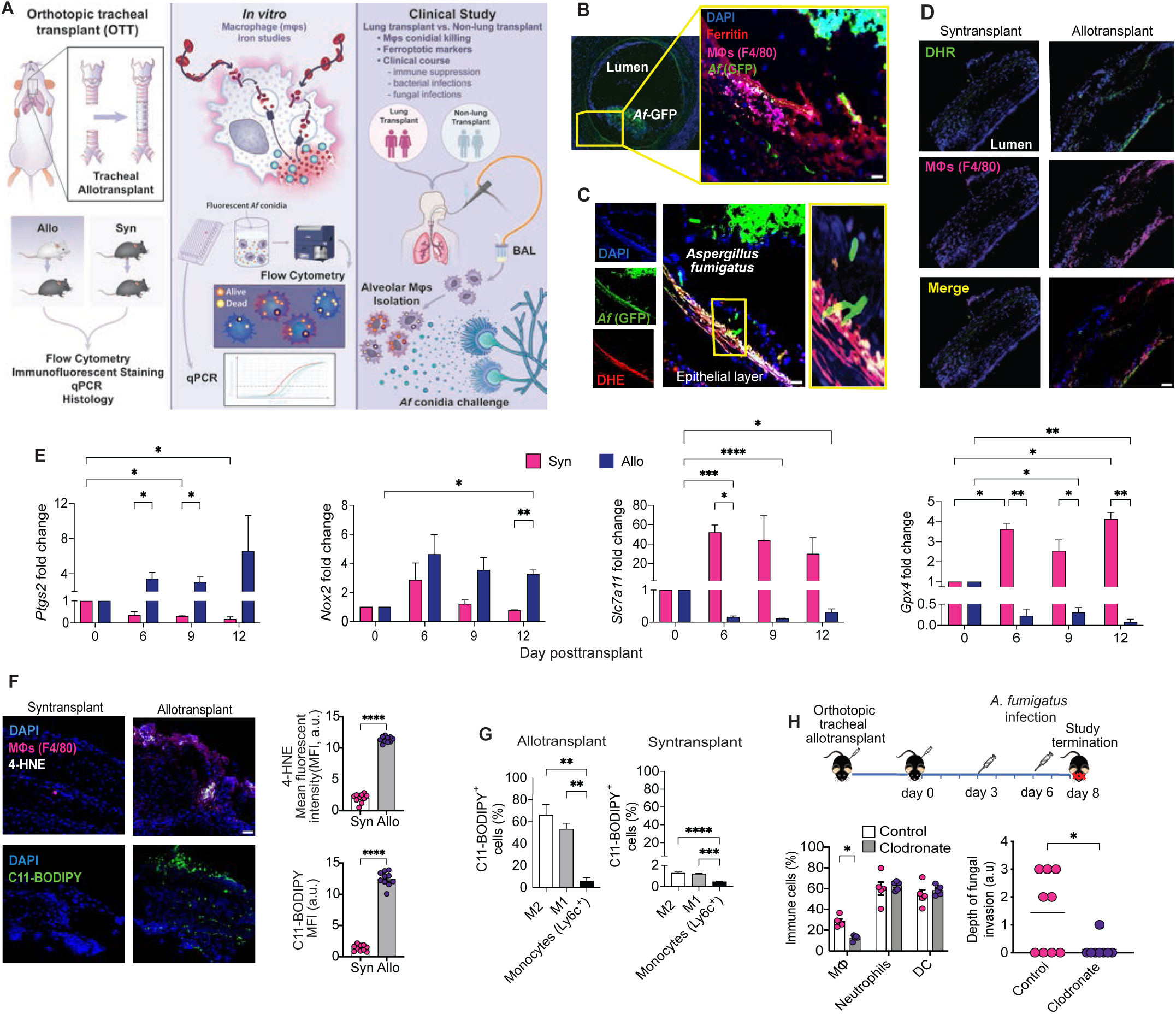
Alloimmune-mediated microhemorrhage increases transplant macrophage ferroptosis. (**A**) Schematic of overall study design. (**B**) Representative immunofluorescent image of explanted allogeneic (allo) tracheal transplant infected with a GFP-labeled *Aspergillus fumigatus* (*Af*-GFP) stained for ferritin (red, protein-bound iron) and macrophages (mфs, F40/80^+^ cells, pink). Scale bar, 50µm. Magnification of image, inset (yellow) (**C**) Representative immunofluorescent image of *Af*-GFP (green) infected allotransplant stained for cell nuclei (DAPI, blue) and dihydroethidium (DHE, red) superoxide detector. Scale bar, 50µm, (**D**) Representative immunofluorescent images of syngeneic (syn) transplant and allotransplant tracheas (day 12 posttransplant), stained for dihydrorhodamine 123 (reactive oxygen species (ROS) detector). Scale bar, 100µm, (**E**) Relative gene expression as measured by real time quantitative polymerase chain reaction (qPCR) of ferroptosis regulators *Ptgs2*, *Nox2, Slc7a11*, and *Gpx4* in allo- and syntransplants by day posttransplant. The linear mRNA of β-actin was used as control. Data analyzed by 2-way ANOVA with Tukey’s test (n = 3/time point) (**F**) Representative immunofluorescent images of syn- and allotransplant tracheas stained for cell nuclei (DAPI), mфs (F4/80) and ferroptosis markers 4-hydroxynoneal (4-HNE, upper panel), and CD11-BODIPY (lower panel). Scale bar, 100µm. Quantification of mean fluorescent intensity (MFI) of ferroptosis markers 4-HNE and C11-BODIPY, as analyzed by t-test (n=10/group). (**G**) Flow cytometry quantification of the percentage of M1 mфs (F4/80^+^, CD11b^+^, CD80^+^, CD86^+^), M2 mфs (F4/80^+^, CD11b^+^, CD206^+^, CD163^+^) and the bone marrow derived monocytes (CD11b^+^, Ly-6C^+^) positive for C11-BODIPY^+^ in allotransplants (left panel) and syntransplants (right panel), as analyzed by one-way ANOVA with Tukey’s test (n=5/group). (**H**) Schematic representation of clodronate mфs depletion experiment (top panel). Allotransplants were treated (every 3 days) with clodronate liposomes while controls were treated with liposomes containing phosphate buffered saline. Animals were infected with *Af* on day 6 posttransplant. Mice were sacrificed 2 days post-infection. Flow cytometry (lower left panel) quantification of percentage mфs (F4/80^+^, CD11b^+^), neutrophils (Ly-6G^+^, CD11b^+^) and dendritic cells (Ly-6C^+^, CD45^+^) in explanted trachea. Depth of fungal invasion measured using a Grocott Methenamine Silver (GMS) stain in explanted trachea with depth of invasion graded (0-4 scale, Supplemental Figure 1A), arbitrary units (a.u.), as analyzed by Mann-Whitney test (n=8-10/group).^15^ For all experiments data presented are mean +/- SEM, \**P* < 0.05, \*\**P* < 0.01, \*\*\**P* < 0.001, ****

## Results

### Orthotopic tracheal transplant

#### Alloimmune-mediated hemorrhage increases transplant oxidative stress

Mφs phagocytose and kill inhaled conidia. Because mφs are central to the defense against *Af* infection and *Af* is a major problem in LTRs, we wanted to understand whether and how mφs are affected by the iron overload observed in the allotransplant. In the OTT model, allotransplants (BALB/c mouse-donor trachea transplanted into a B6 mouse recipient) undergo graft rejection, as no immune suppression is given, resulting in rejection-mediated microvascular damage, hemorrhage, and iron overload.(14) In contrast, syngeneic transplants (B6 donor and B6 recipient) do not reject the transplant and do not experience iron overload. Allotransplants were infected with an *Af* strain (*Af293*) expressing green fluorescent protein (*Af-*GFP) and excised tracheas were stained with antibodies against ferritin (iron) and mφs. Widespread colocalization of mφs, and intracellular and extracellular ferritin, was seen at the site of *Af* infection (Figure 1B). As iron overload can increase the oxidative stress of the microenvironment through the Fenton reaction, the level of ROS in tissues was evaluated in the allotransplant.(25) *Af*-GFP infected allotransplants were stained with the superoxide detector dihydroethidium (DHE). Fluorescence microscopy identified high levels of ROS at sites of *Af* invasion (Figure 1C). To clarify ROS production specifically from mφs, tracheal grafts from allo- and syntransplants were stained with dihydrorhodamine 123 (DHR), a ROS detector, and antibodies to murine mφs. In allotransplants, mφs demonstrated high levels of ROS, a finding not seen in syntransplants (Figure 1D).

#### Alloimmune-mediated macrophage ferroptosis promotes fungal invasion

Having demonstrated iron overload and increased ROS in the OTT, we posited that the increased alloimmune-mediated oxidative stress was due to ferroptosis. Using quantitative real-time polymerase chain reaction (qPCR), we first investigated in non-infected, explanted trachea the expression of ferroptosis regulators prostaglandin endoperoxide synthase 2 (*Ptgs2*), nicotinamide adenine dinucleotide phosphate (NADPH) oxidase 2 (*Nox2*), *Gpx4* and solute carrier family 7 member 11 (*Slc7a11*) in allotransplants and syntransplant over time. *Slc7a11* is a gene that governs cysteine transport, a critical component in maintaining cellular redox levels.(26) Expressions of *Ptgs2* and *Nox*2 were increased in the allotransplants relative to syntransplants and controls after 6 days posttransplant. Expression of the anti-ferroptotic genes, *Gpx4,* and *Slc7a11*, were decreased relative to levels in non-transplant controls and syntransplants (Figure 1E). To verify ferroptosis in transplanted tissues, allo- and syntransplants were stained with antibodies against mφs and 4-hydroxynoneal (4-HNE), a product of lipid peroxidation, and examined using fluorescence microscopy. High levels of 4-HNE were observed in mφs from allo-but not syntransplants (Figure 1F). Mφ lipid peroxidation was further investigated by tissue staining and flow cytometry, using C11-BODIPY, a detector of lipid peroxidation (Figure 1F). Allotransplants showed a significant increase in ferroptotic mφs compared to syntransplants (mean percentages of C11-BODIPY^+^ mφs: 49.2% and 1.7%, respectively, *P* = 0.003). To better understand the impact of ferroptosis on mφ subpopulations, C11 BODIPY^+^ mφs isolated from explanted trachea were gated to differentiate between classically activated (M1) and alternatively activated (M2) subpopulations, as well as mφs from bone marrow derived monocytes (i.e., transplant recipient). Flow cytometry results showed that M1 and M2 mφs had significantly higher degrees of lipid peroxidation than ones derived from pro-inflammatory monocytes (Figure 1G). Collectively, these findings suggest that transplant mφs undergo ferroptosis and that recruited mφs are polarized towards a pro-inflammatory, more ferroptosis-resistant subpopulation.(22)

To investigate whether these iron-overloaded mφs may paradoxically increase the risk of *Af* invasion, mouse tracheal transplant recipients were treated with clodronate liposomes to deplete mφs (3 days before transplant and every 3 days after transplant, see schematic Figure 1H). Allotransplants treated with liposomes containing phosphate-buffered saline (PBS) were studied in parallel as controls. Flow cytometry was used to verify the efficacy of mφ depletion and to assess the effect of clodronate treatment on other professional phagocytes (neutrophils and dendritic cells). Clodronate treatment effectively decreased OTT mφs compared to control animals (12.8% and 28.5%, respectively), whereas the proportions of neutrophils and dendritic cells were similar in both groups (Figure 1H). Having demonstrated the effectiveness of mφ depletion, we sought to investigate the degree of *Af* invasion in clodronate liposome-treated transplants compared to recipients treated with PBS-loaded liposomes. Transplants were infected on day 6 and euthanized 2 days post-infection. The depth of fungal invasion was graded histologically using Grocott’s Methenamine Silver (GMS) staining for *Af* in the explanted OTT sections (Supplemental Figure 1A).(12–14) *Af* invasion in the OTT was significantly reduced in mφ-depleted mice compared to controls (Figure 1H, Supplemental Figure 1B).

#### Ferroptotic stress attenuates macrophage conidial killing through a loss of phagolysosomal acidification

To examine the impact of exogenous iron overload, primary mφs were isolated from mouse lungs and exposed in culture to increasing iron concentrations (at levels consistent with tissue microhemorrhage and a mouse hematocrit of 20% to 60%).(14, 27) Cell viability was measured using the Sulforhodamine Beta (SRB) assay. Primary mφs in complete media (control), as well as those treated with iron and/or the iron chelator deferasirox (DFX), were studied in parallel. Iron induced a dose-dependent decrease in viable mφs compared to control and DFX treated mφs (Figure 2A).

**Figure 2.**
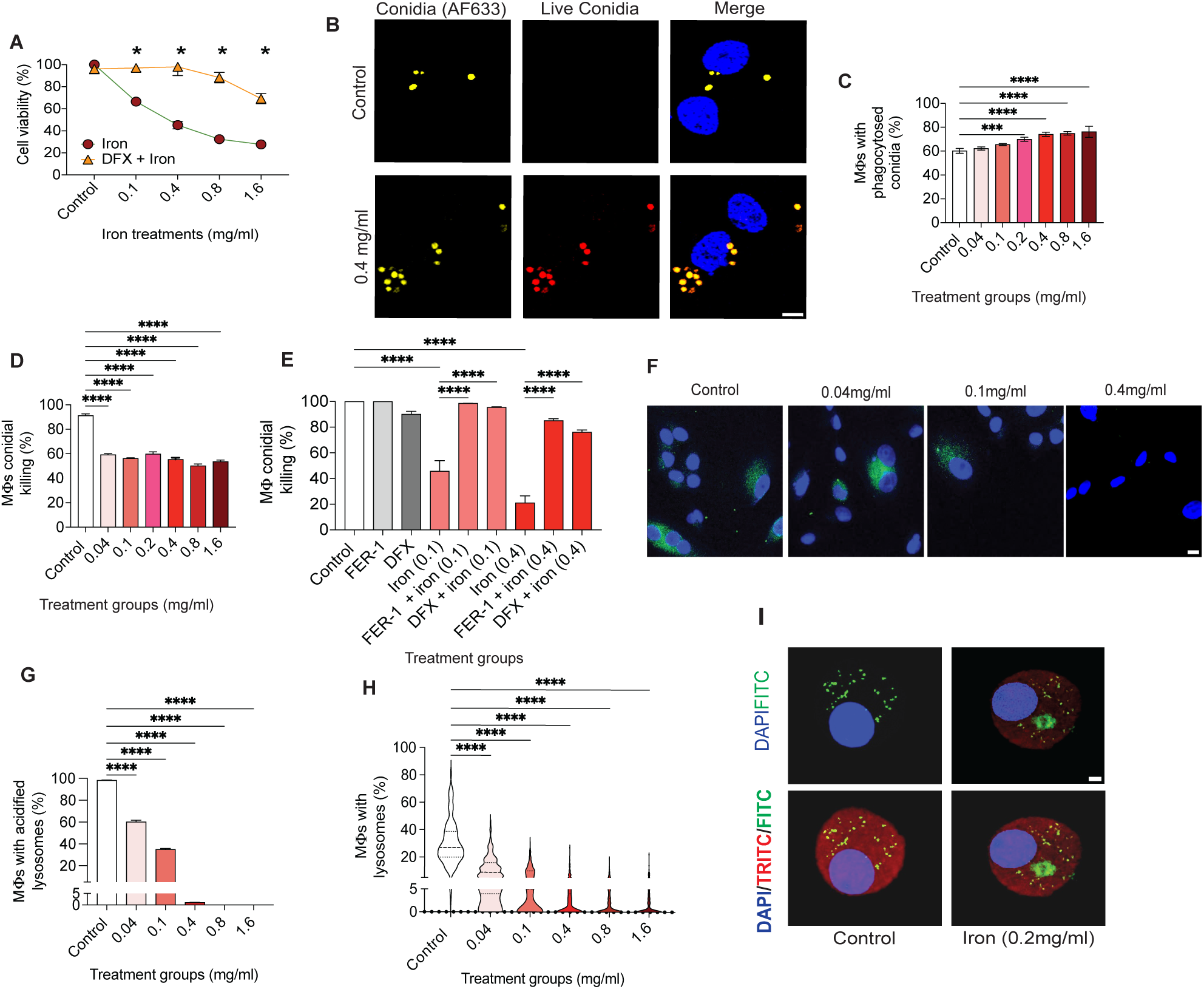
Ferroptotic stress attenuates macrophage conidial killing through loss of phagolysosomal acidification. (**A**) Lung mф cell viability, as measured by Sulforhodamine Beta (SRB) assay, comparing primary lung mфs in complete media alone (control) and those treated with increasing iron concentrations (0.1, 0.4, 0.8, 1.6 mg/ml) alone or with deferasirox (DFX 100µM, an iron chelator), analyzed by one-way ANOVA (n = 3/group). (**B**) Representative immunofluorescent images of primary mouse lung mфs co-cultured with the *Af*-<u>fl</u>uorescent *<u>A</u>spergillus* <u>re</u>porter (*Af*-FLARE) conidia (1:5 ratio) for 6 hours. Live conidia have both fluorescent markers (orange, composed of endogenously expressed DsRed (red) and cell wall coating with AF633 (yellow), merged image bottom right panel), whereas killed conidia have only the cell wall tracer (yellow, merged image upper panel) Representative fluorescent images of *Af*-FLARE conidial killing in controls (upper panels) and mф exposed to 0.4 mg/ml iron (lower panels). Scale bar, 5µm (**C**) Flow cytometry analysis of primary mф exposed to increasing iron conditions for 16 hours (0.04, 0.1, 0.2, 0.4, 0.8, 1.6 mg/ml). Percentage conidial phagocytosis and **(D)** *Af*-FLARE conidial killing, calculated by percentage phagocytosed live conidia/total phagocytosed conidia (live and dead conidia). Data analyzed by one-way ANOVA (for phagocytosis experiment n = 3/group, for conidial killing experiments n = 5/group). (**E**) Flow cytometry analysis of percentage conidial killing for primary mфs cultured with *Af*-FLARE conidia, iron (0.1 and 0.4 mg/ml) and/or with FER-1 (ferroptosis inhibitor, 10 μM) and deferasirox (DFX, 20 μM) (6 hours) analyzed by one-way ANOVA (n=3/group). (**F**) Representative images of lysosomal acidification using LysoTracker Green DND-26. Primary mф exposed to increasing iron concentrations (0.04, 0.1, 0.4 mg/ml) and lysosomal acidification indicated by LysoTracker (green) fluorescence. Cell nuclei were stained using DAPI (blue). Scale bar, 10µm. (**G**) Quantification of the percentage of acidified lysosomes in primary mфs cultured with increasing iron concentrations (0.04, 0.1, 0.4, 0.8 1.6 mg/ml) or media alone (control). Data analyzed by one-way ANOVA, (n = 3 biologic and technical replicates/group). (**H**) Violin plot depicting the percentage of mфs with lysosomes, as measured by lysosomal-associated membrane protein (LAMP)-1 staining. Mфs cultured with increasing iron concentrations (0.04, 0.1, 0.4, 0.8 1.6 mg/ml) or media alone (control). Data analyzed by one-way ANOVA, (n = 3 biologic and technical replicates/group). (**I**) Representative fluorescence images of lysosomal leakage for primary mфs cultured in complete media alone (control, left panel) and iron dextran (0.2 mg/ml, right panel) and stained with TRITC-dextran (red)/FITC-dextran (green) and DAPI (cell nuclei, blue). Scale bar, 5µm. Data presented are mean +/- SEM \**P* < 0.05, \*\**P* < 0.01, \*\*\**P* < 0.001, \*\*\*\**P* < 0.0001.

We then examined the impact of mφ ferroptotic stress on phagocytosis and killing of *Af* conidia. Primary mφs isolated from mouse lungs were co-cultured with the *Af*-<u>fl</u>uorescent *<u>A</u>spergillus* <u>re</u>porter (*Af*-FLARE) conidia. Live conidia have both fluorescent markers (orange, composed of endogenously expressed DsRed (red) and cell wall coating with AF633 (yellow)), whereas killed conidia have only the cell wall tracer (yellow) (Figure 2B).(28, 29) We found that increasing iron concentrations (>0.2 mg/ml) slightly increased conidial phagocytosis (Figure 2C) but substantially decreased killing of phagocytosed *Af-*conidia (Figure 2D). Conidial killing was then assessed in mφs cultured with iron (0.1, 0.4 mg/ml) alone or co-cultured with FER-1 (10μM) or DFX (20μM). Ferrostatin-1 and DFX treatment restored mφ-mediated *Af-*conidial killing at 0.1 mg/ml iron concentrations, an effect that persisted but to a lesser degree at an iron concentration of 0.4mg/ml (Figure 2E). Mφs ferroptosis was verified using C11 BODIPY staining, which confirmed that FER-1 and DFX significantly reduced the ferroptotic stress in primary murine mφs (Supplemental Figure 2A). Taken together, these data strongly suggest that ferroptosis impairs the mφs’ ability to kill phagocytosed *Af* conidia and that anti-ferroptotic treatments can rescue fungal killing.

To enhance insight into the cellular mechanism underlying the decrease in conidial killing, we investigated the impact of iron overload on lysosomal acidification, a necessary step for *Af* enzymatic killing occurring in the phagolysosome.(30) Lysosomal acidification, as measured by LysoTracker staining, demonstrated an iron dose-dependent decrease in the number of cells with acidified lysosomes (Figure 2F, and G).(31) Lysosomal loss was confirmed by lysosomal-associated membrane protein (LAMP)-1 staining (Figure 2H). Iron caused a significant pH increase in mφs, as measured by the ratio of LysoSensor yellow to blue dextran fluorescent probes (Supplemental Figure 2B).(32) To investigate whether lysosomal leakage is involved in the loss of acidified lysosomes, primary mouse lung mφs were treated with increasing iron concentrations, stained with a TRITC-dextran/FITC-dextran, and examined by fluorescence microscopy.(33) Extensive lysosomal leakage was observed at iron concentrations > 0.2 mg/ml (Figure 2I). Together these results indicate that iron impairs phagolysosomal acidification through lysosomal leakage and loss.

### Ferroptosis studies in human lung transplant recipients

#### Ferroptotic stress is associated with fungal infections in lung transplants

The clinical significance of our findings was investigated in bronchoscopy samples from a convenience sample obtained in the usual clinical care of adult LTRs (n=15) and non-LTR controls (n=11). Three controls samples (27%, 3/11) were obtained from patients on immune suppression after heart transplantation (Supplementary Table 1 for full details of study participants, see Methods for inclusion and exclusion criteria and study procedures). Iron content and ferroptosis markers of lipid peroxidation (4-HNE, lipid hydroperoxide (LPO), and malondialdehyde (MDA)) were assayed in BAL supernatant. All makers of ferroptosis were significantly higher in LTRs than controls (Figure 3A). Alveolar mφs were isolated from the BAL pellet and co-cultured with *Af-*FLARE conidia for 6 hours. We evaluated the ability of patient-derived mφs to phagocytose and kill *Af* conidia by flow cytometry. Alveolar mφs isolated from LTRs were defective in both conidial phagocytosis (Figure 3B) and killing (Figure 3C) compared to mφs isolated from non-LTRs. Clinical information including age, sex, immune suppressive medications and microbiologic results was also correlated with markers of ferroptosis (4-HNE, MDA, LPO) and iron. Here, we present the correlation coefficient (r value) for both LTRs and non-LTRs (Figure 3D). There was a strong correlation between increased 4-HNE and to a lesser extent MDA and LPO with decreased mφ conidial killing. Iron content of the BAL fluid was moderately correlated with conidial killing. Markers of ferroptosis also were significantly associated with any clinical history of proven or probable fungal infection as per European Organization of Research and Treatment of Cancer-Mycoses study group (EORTC-MSG) guidelines (Figure 3E and F).(34)

**Figure 3.**
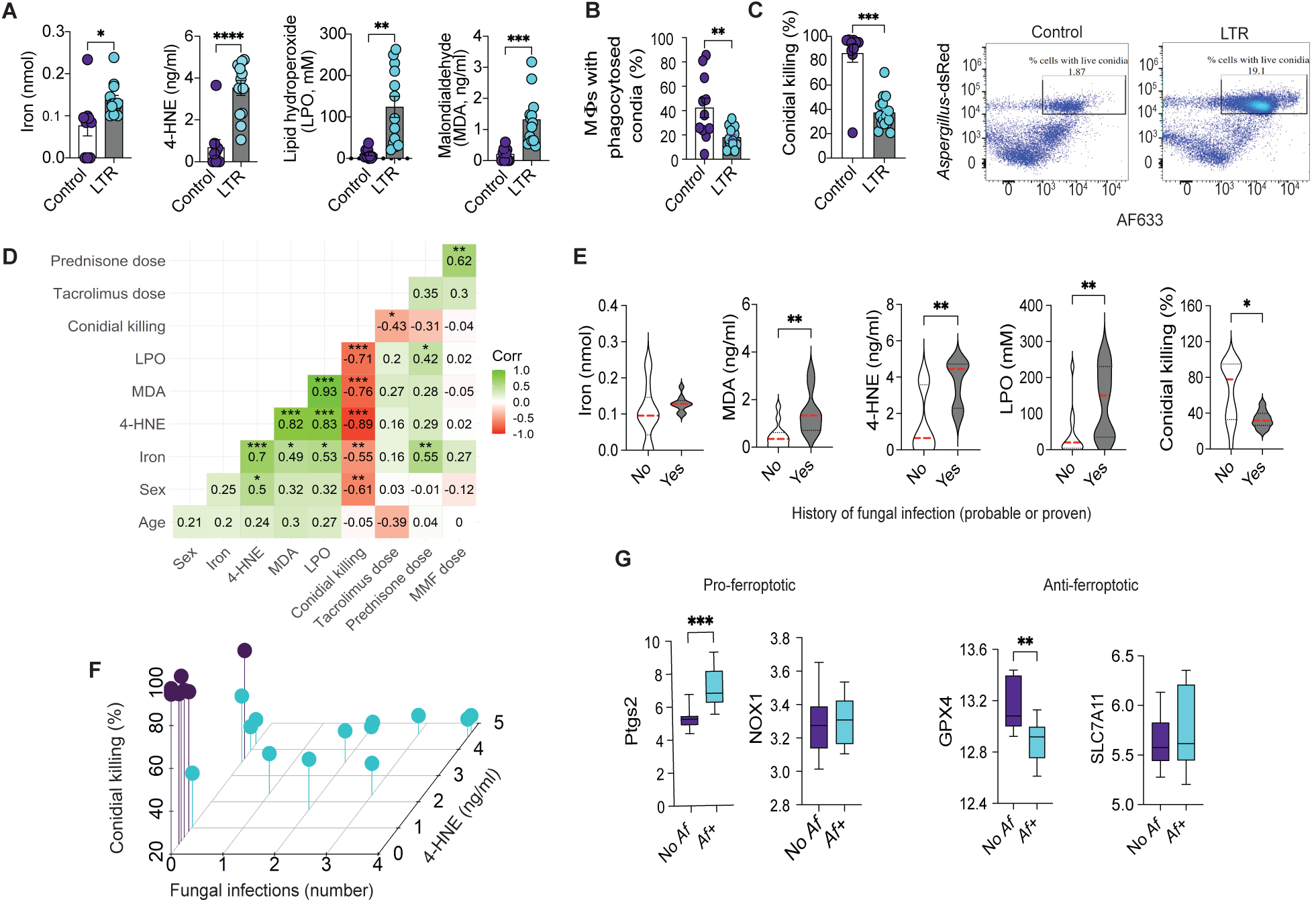
Ferroptotic stress is associated with decreased conidial killing and a history of clinical infections. (**A**) Bronchoalveolar lavage (BAL) fluid samples were collected from lung transplant recipients (LTR, n=15) and non-lung transplant (controls, n=11) and analyzed for iron (nmol), 4-HNE (ng/ml), lipid hydroperoxide (LPO, mM) and malondialdehyde (MDA, ng/ml) content. (**B**) Flow cytometry analysis of alveolar mfs isolated from patient BAL samples and co-cultured with *Af*-FLARE conidia (6h) (ratio 1:5). Quantification of mfs percentage with phagocytosed conidia. (**C**) Quantification of mfs percentage *Af* conidial killing (live/total (dead+ live) phagocytosed conidia) in LTR and controls (left panel). Representative flow cytometry dot blots (right panels). (**D**) Correlation matrix analysis comparing markers of ferroptosis (iron, 4-HNE, LPO, MDA) with conidial killing, age, gender and clinical immune suppression. (**E**) Association between iron and markers of ferroptosis and a clinical diagnosis of proven or probable fungal infections by EORTC/MSG guidelines.^38^ Analyzed by Mann Whitney U test. (**F)** Association between conidial killing, 4-HNE and number of fungal infections (top panel) and number of acute or chronic bacterial infections (bottom panel). (**G**) Gene set variance analysis of LTR colonized with *Af* (*Af+*, n=12) and LTR without evidence of *Af* colonization (No *Af*, n=10) accessed from ebi.ac.uk (36) to generate gene set enrichment scores for *PTGS2, NOX1, GPX4*, and *SLC7A11*). Analyzed using the Mann-Whitney U test. Data presented are median and high and low. Unless otherwise noted all other data presented are mean +/- SEM \**P* < 0.05, \*\**P* < 0.01, \*\*\**P* < 0.001, \*\*\*\**P* < 0.0001.

To analyze the association of ferroptosis markers with mφ conidial killing and a clinical history of fungal infection, we generated univariate and multivariate regression models controlling for clinical factors including age and commonly used immune suppressive medications (i.e., tacrolimus, prednisone, and mycophenolate mofetil (MMF)). In univariate analyses, all ferroptosis markers showed a significant inverse relationship with conidial killing. For fungal infection, only MDA, 4-HNE, and LPO had significant associations (Supplementary Table 2). The relationship between mφ conidial killing, ferroptosis markers and iron was maintained in multivariate models adjusting for age and the daily dose of immune suppressive medications (Supplementary Table 3, 4 and 5). Together these data support the interpretation that alloimmune-mediated iron overload induces alveolar mφ ferroptosis and an inability to kill *Af* conidia.

#### Lung transplant BAL transcriptome is characterized by lipid metabolism and regulators of ferroptosis

The transcriptome of the whole human BAL was evaluated using bulk RNA-sequencing (RNA-seq). RNA was extracted and library preparation was performed, using a previously published pipeline optimized for BAL.(35) Non protein-coding, mitochondrial, and ribosomal reads were excluded and resulting protein-coding transcript counts underwent gene set variance analysis (GSVA) using Reactome and Gene Ontology: Biological Function repositories to produce semi-normally distributed gene set enrichment scores. Gene set enrichment scores were compared between LTRs (n=6) and controls (n=5) and correlated with BAL measures of conidial killing and markers of ferroptosis. The expression of the BAL Reactome gene sets “Autophagy of peroxisomes” and “Peroxisomal lipid metabolism” were non-significantly increased in LTRs (Supplemental Figure 3A). The expression of the “Autophagy of peroxisomes” gene set was moderately correlated with MDA (r = 0.75, *P* = 0.008), LPO (r = 0.73, *P* = 0.015), and 4-HNE (r = 0.67, *P* = 0.027) and inversely correlated with conidial killing (r = −0.69, p = 0.023). Expression of the “Peroxisomal lipid metabolism” gene set was moderately correlated with MDA (r = 0.6, *P* = 0.05) and LPO (r = 0.6, *P* = 0.05). To validate these preliminary transcriptome findings, data from the “Gene Expression Profiling of Bronchoalveolar Lavage Cells During *Aspergillus* Colonization of the Lung Allograft” study were accessed via ebi.ac.uk (E-MTAB-6040).(36) As previously reported, this dataset includes adult LTRs, of whom 55% (12/22) had BAL with *Aspergillus* colonization. Normalized transcript abundance was compared for genes of interest (*PTGS2, GPX4, NOX1*) and GSVA was applied, as above, to generate gene set enrichment scores to the Reactome collection. We showed a similar non-significant increase in expression of the “Autophagy of peroxisomes” gene set, whereas expression of the “Peroxisomal lipid metabolism” gene set was similar between the two groups (Supplemental Figure 3B). For individual gene expression studies, there was a significant increase in the expression of *PTGS2* (*P* = 0.001) and a decrease in the expression of *GPX4* (*P* = 0.004) in LTRs colonized with *Af* compared to LTRs without fungal colonization (Figure 3G). In LTRs colonized with *Af*, the positive ferroptosis regulator *NOX1* and the negative regulator *SLC7A11* trended toward a non-significant increased and decreased expression, respectively. Functional enrichment analysis of the upregulated genes from the LTR with *Aspergillus* colonization revealed that several of the over-represented (*P* < 0.05) biological processes were relevant to iron metabolism, ferroptosis and ROS, i.e., iron ion transport (GO:0006826), transition metal ion transport (GO:0000041), response to lipid (GO:0033993), response to oxygen-containing compound (GO:1901700), programmed cell death (GO:0012501), and cell death (GO:0008219) (Supplementary Table 6).(36) Pathway analysis indicated that iron-overload related pathway ‘defective SLC40A1 causes hemochromatosis 4 (macrophages) (R-HSA-5619049)’ was identified as one of the most over-represented pathways (fold change: 52.7, *P* = 0.000358, false discovery rate: 0.049).(36) Together, the results from the human BAL transcriptome and the analysis of gene sets from the published literature are consistent with our findings, suggesting that ferroptosis plays an important role in fungal clearance in the LTR.

## Discussion

Currently, the focus of treating infections with *Af,* a ubiquitous and emerging pathogen, in transplant recipients is anti-fungal therapy and modulation of recipient immune function. Our study is the first to identify mφ ferroptosis occurring after lung transplantation as a risk factor for host susceptibility to fungal infections. Our results suggest that ferroptosis impacts mφ killing through a decreased lysosomal acidification and lysosomal loss, facilitating *Af* invasion. Multiple aspects of this study support this conclusion. First, our investigation of ferroptosis hallmarks in the mouse OTT model of invasive aspergillosis demonstrated that mφs in the allotransplant graft predominantly underwent ferroptosis. *In vitro* studies revealed that ferroptotic mφs cannot effectively kill *Af* conidia. Results from the OTT model and *in vitro* findings were corroborated by clinical data from human alveolar mφs and BAL, comparing lung transplant and non-lung transplant patients, and in published gene sets from LTRs with a history of *Af* colonization.

Ferroptosis is characterized by an iron-dependent increase in ROS because of lipid peroxidation, causing non-enzymatic damage that affects the integrity of the cellular membranes and alters their fluidity and permeability.(37) Ferroptosis has been described in lung and renal transplantation in the setting of ischemia reperfusion injury.(38, 39) To our knowledge, we are the first to describe a persistent ferroptotic state in lung transplants. In prior animal studies, we showed that transplant graft iron overload stems from rejection-mediated vascular damage and tissue bleeding. Here, we extend these findings to show that the mφ erythrophagocytosis required to process iron overload from hemorrhage in the allograft results in the inability of mφs to effectively acidify phagolysosomes. These findings corroborate those from a study by Kao et al., showing that the monocyte and mφ cell line (THP-1) exposed to iron had a decrease in lysosomal acidification.(40) This iron-induced loss of lysosomal acidification suppresses the hosts’ capacity to eliminate phagocytosed pathogens. We suspect that the profound loss of lysosomal acidification observed at relatively low iron levels (0.04mg/ml) may explain in part a lack of a dose-dependent effect of iron on conidial killing. These findings are clinically significant, as anti-ferroptotic therapies (FER-1, or DFX) restored conidial killing. Notably, alveolar mφs isolated from LTRs were defective in conidial phagocytosis, whereas this was not observed in our *in vitro* primary murine mφ studies. Possible differences between these findings may be related to exogenous immune suppressive agents used in solid organ transplants such as MMF, and prednisone which can impact monocyte and mφ phagocytosis and are commonly used in LTRs.(41, 42)

Pathogens have been shown to manipulate ferroptotic pathways to enhance their survival and propagation in the host. Infection with Hepatitis B virus results in liver cirrhosis in part by downregulating *SLC7A11* expression, causing liver damage through hepatocyte ferroptosis.(26) In contrast, the oncogenic pathogen, Epstein Barr virus (EBV), increases *GPX4* expression through downregulation of nuclear factor erythroid 2-related factor, protecting against ferroptosis and increasing resistance of EBV-infected nasopharyngeal cancer cells to chemotherapeutic drugs.(43) Single-cell RNA-seq studies have suggested that infection with severe acute respiratory syndrome coronavirus-2 (SARS-CoV2) upregulates lymphocyte expression of ferroptosis-related genes and viral infection has been implicated in cardiac sinoatrial node cell ferroptosis.(26) Recently, *Mycobacterium tuberculosis* tissue invasion and disease dissemination have been shown to be promoted by induction of mφ ferroptosis through downregulation of *GPX4* expression.(44) Indeed, FER-1 treatment of mice infected with *M. tuberculosis* increased bacterial clearance.(45) Among fungal pathogens, *Magnaporthe oryzae*, which is responsible for destructive infections in cultivated rice crops, manipulates mitochondrial oxidase, increasing lipid ROS and cell death in infected rice cells.(26) *Cryptococcus neoformans*, the causative agent of meningitis in persons with the acquired immune deficiency syndrome, increases levels of lipid ROS and depletes brain glutamate, resulting in ferroptosis.(26) In contrast to these examples, our data demonstrate that host immune cell ferroptosis can promote immune dysregulation and establishment of infection. Interestingly, aside from the postulated association for ferroptosis in the setting of concomitant viral infections (see below), the relationship between infection with *Mucorales* species, iron avid molds and ferroptosis has not been previously described.

Limitations to the current study require mention. First, ferroptosis is not cell dependent; it can occur in a variety of cells, including epithelial cells, endothelial cells, neutrophils, dendritic cells, and lymphocytes.(46) Since tissue resident mφs are the first line of defense against fungal pathogens, we focused our investigation on the impact of ferroptosis on this innate immune cell population. Additional research is needed to examine whether and how lung transplant-related ferroptosis may affect other cell populations and cellular interactions involved in the host defense against pathogens. Second, we did not explore a potential additive role of *Af* virulence factors in promoting ferroptosis. Recent studies have examined the *Af* secondary metabolite, gliotoxin, for its ferroptosis-inducing capacity in lung adenocarcinoma and breast cancer cell lines.(47) Thus, a pro-ferroptotic role for *Af* may be posited, as has been shown with other pathogens. Ongoing studies in our lab are examining the pro-ferroptotic impact of *Af* infection in transplant and non-transplant mouse models. One might also hypothesize that ferroptosis could play a role in the concomitant fungal and viral infections in persons infected with SARS-CoV2. A posited ferroptosis-based association of coinfecting pathogens may explain not only the association of *Af* and SAR-CoV2, but also the higher prevalence of infection with *Mucor* species and SARS-CoV2.(48, 49) In persons with COVID-19 associated pulmonary mucormycosis, iron-metabolism and ferroptosis genes were upregulated in innate immune cells compared to healthy controls.(50) Finally, while the current study found a strong association between iron overload and markers of ferroptosis other blood proteins could have contributed to a ferroptotic microenvironment. In our previous study we found that both iron and hemoglobin can promote *Af* invasion.(14)

In conclusion, this study identifies mφ ferroptosis as a novel form of immune dysfunction. We have previously shown a determinant role of transplant iron overload due to microvascular damage and hemorrhage in *Af* invasion of the graft.(14) Here we extend these findings to show a mechanistic and clinical link between allograft microvascular rejection and fungal invasion through an inability of lung transplant mφs to eliminate phagocytosed *Af* conidia. Moreover, we show that iron overload impairs mφ lysosomal acidification and that anti-ferroptotic treatments can rescue clearance. As increased iron has been implicated in multiple pulmonary diseases, including chronic obstructive pulmonary disease, asthma, cystic fibrosis and the acute respiratory distress syndrome (conditions also characterized by a higher incidence of *Af*-related pulmonary diseases), our findings may have more far-reaching implications.(51) Future studies should also consider the impact of ferroptosis on infections in hematopoietic cell transplant recipients who are also characterized by high iron levels and a high incidence of *Af* infection.(52) Anti-ferroptotic studies in lung transplants and other pulmonary diseases would be impactful, as mitigation of ferroptosis through FER-1 and DFX treatment may offer novel strategies to improve *Af* clearance.

## Methods

### Sex as a biological variable

Our study exclusively examined male mice. It is unknown whether the findings are relevant for female mice. For human studies, our study examined males and females, and similar findings are reported for both sexes.

### Study approval

Animal experiments were approved by Stanford’s Laboratory Animal Care (APLAC 33935).. The human study was approved by the Institutional Review Board of Stanford University (IRB #59128). All the patients or legally authorized representatives signed consent forms. Bronchoalveolar Lavage (BAL) samples were collected from patients at Stanford University Hospital. Stanford’s Administrative Panel on Biosafety (APB 3959) approved all *Aspergillus fumigatus (Af*) experiments.

### Aspergillus fumigatus culture and preparation of inoculum

Dr. David Stevens provided the *Af*293-GFP and 10AF strains. Dr. Tobias Hohl provided the *Af*293 DsRed FLARE strain which was labeled with AF633 tagging as previously described.(28, 29) *Af* conidia cultures were prepared as previously described.(12, 14) For *in vitro* assays conidial suspensions were 10^5^ conidia/ml and for *in vivo* experiments 10^8^ conidia/ml were used.(12, 14)

### Orthotopic tracheal transplantation (OTT) Af infection model

Five-week-old male C57BL/6, and BALB/c, performed, as previously described.(12–14) Briefly, mice were inoculated intratracheally with *Af* conidia (40 μl). Depth of *Af* infection in infected explanted tracheas was graded using Grocott’s methenamine silver staining (0-4 semi-quantitative scale, Supplemental Figure 1A), as previously described.(12–14)

### Studies in OTT explanted tissues

#### Tracheal tissue immunostaining

Frozen tissues were sliced (5 µm) using a Cryostat (Leica CM1860 UV). Sections were fixed and permeabilized with 4% paraformaldehyde and 0.1% Triton X-100 respectively. Fungal elements were stained using calcofluor white (Sigma-Aldrich #18909) or Fluorescent Brightener 28 disodium salt solution (Sigma-Aldrich #910090) supplemented by 10% potassium hydroxide (Ricca Chemical #61304). Sections were incubated with blocking solution (1% BSA, 22.52 mg/mL glycine in PBST) for 30 min and then incubated with primary antibodies anti-4-HNE (Abcam #ab48506), anti-COX2 (Santa-Cruz biotechnology #sc-376861 AF647), anti-ACSL4 (Abcam # ab240135), anti-F4/80 (Abcam #ab16911), anti-ferritin (Abcam #ab75973) in blocking solution for 1h at RT. When secondary Abs were needed sections were incubated with appropriate fluorophore-conjugated secondary antibodies (goat anti-rabbit IgG H&L Abcam #ab150077), goat anti-rat IgG H&L Abcam #ab150157), for 1h at RT in the dark. Slides were mounted using Invitrogen™ ProLong™ Gold Antifade Mountant with DAPI. Images and z stacks were taken using a confocal microscope (Leica TCS SP8, Leica STELLARIS).

### C11 BODIPY studies

#### Imaging

Frozen tracheal sections (5µm) were stained with 5mM C11 BODIPY (Invitrogen #D3861) for 30 min in a 5% CO2 atmosphere at 37 °C. Sections were washed with PBS and fixed with 4% PFA for 10 min at RT. Sections were washed with PBS and mounted using Invitrogen™ ProLong™ Gold Antifade Mountant with DAPI. Confocal microscopy images were obtained using confocal microscope (Leica TCS SP8, Leica STELLARIS).

#### Flow Cytometry

Single cell suspensions from mouse trachea homogenates were generated. Cells were incubated with 1.5 µM C11-BODIPY 581/591 (Invitrogen, cat. no. D3861) for 30 min in a 5% CO2 atmosphere at 37 °C. Subsequently, cells were washed with PBS. Nonspecific staining sites were blocked with Fc block anti-mouse CD16/CD32 BD Pharmigen #553142) for 15 min at 4 °C. Cells were washed with PBS, 2% FBS, 0.5 mM EDTA, and stained with anti-mouse F4/80 AF488 BioLegend #123120, anti-mouse CD11b PE/Dazzle 594 BioLegend #101256, anti-mouse CD80 BV711 BioLegend #104743, anti-mouse CD86 BV785 BioLegend #105043, anti-mouse CD206 AF647 BioLegend #141712, and anti-mouse CD163 BV421 BioLegend #155309. Cells were washed with PBS, 2% FBS, 0.5 mM EDTA, passed through a 70μm cell strainer and analyzed with a flow cytometer (BD® LSR II, Agilent NovoCyte Quanteon, Agilent NovoCyte Penteon, BD® FACSymphonyTM). Flow cytometry data were analyzed using Flowjo (BD).

### Cell lines

#### Determining cell phenotype using flow cytometry

Mouse lung mφs were isolated, as previously described.(30). *Mφ subpopulations.* Nonspecific staining sites in isolated murine mφs were blocked with Fc block anti-mouse CD16/CD32 BD Pharmigen #553142) for 15 min at 4 °C. Cells were washed with PBS, 2% FBS, 0.5 mM EDTA and stained with anti-mouse F4/80 AF488 BioLegend #123120, anti-mouse CD11b PE/Dazzle 594 BioLegend #101256, anti-mouse CD80 BV711 BioLegend #104743, anti-mouse CD86 BV785 BioLegend #105043, anti-mouse CD206 AF647 BioLegend #141712, and anti-mouse CD163 BV421 BioLegend #155309.

#### Immunophenotyping (mφs, neutrophils, dendritic cells)

Nonspecific staining sites in single cell suspensions from mouse trachea homogenates were blocked with Fc block anti-mouse CD16/CD32 BD Pharmigen #553142) for 15 min at 4°C. Cells were washed and stained with anti-mouse F4/80 AF488 BioLegend #123120, anti-mouse CD11b PE/Dazzle 594 BioLegend #101256, anti-mouse CD11b PE/Dazzle 594 BioLegend #101256, anti-mouse Ly-6G BV570 BioLegend #127629, anti-mouse CD45 BV785 BioLegend #103149, and anti-mouse Ly-6C BioLegend #128037. Cells were analyzed using an LSRII flow cytometer (BD) and data were analyzed using Flowjo (BD).

### Conidial phagocytosis and conidial killing studies

#### Imaging

Mouse lung mφs were plated in sterile coverslips. Mφs were allowed to adhere by incubating at 37° C with 5% CO2 overnight. Iron-dextran (Sigma #D8517) treatments were added for 16h. Lysosomes were stained with CellLight Reagent *BacMAn 2.0* (Invitrogen #C10596) according to the manufacturer’s instructions. 10^5^ FLARE AF633-tagged conidia were added to each well and samples were incubated for 6h at 37° C with 5% CO2. Cells were thoroughly washed with PBS, fixed and permeabilized using 4% paraformaldehyde and 0.1% Triton X-100 respectively. Samples were mounted using Invitrogen™ ProLong™ Gold Antifade Mountant with DAPI. Images and z stacks were taken using a confocal microscope (Leica TCS SP8 and Leica STELLARIS).

#### Flow Cytometry

Mouse lung mφs were plated in 6 well plates and allowed to adhere by incubating at 37° C with 5% CO2 overnight. Iron dextran treatments were added for 16h. FLARE AF633-tagged conidia/ml (10^5^) were added to each flask and samples were incubated for 6h at 37° C with 5% CO_2_. Cells were detached using mφs detachment solution DXF (PromoCell #C-41330). For FER-1 and DFX studies primary murine mφs were cultured with iron dextran (0.1 or 0.4mg/ml) alone or with FER-1 (10μM) or DFX (20 μM) for 16h. FLARE AF633-tagged conidia/ml (10^5^) were added to each flask and samples were incubated for 6h at 37° C with 5% CO_2_. Cells were stained using F4/80 antibody (Invitrogen #MF48021) for 1h at 4° C. Cells were washed 3 times with PBS and analyzed immediately by flow cytometry (BD^®^ LSR II, Agilent NovoCyte Quanteon, Agilent NovoCyte Penteon, BD^®^ FACSymphony^TM^).

### Reactive oxygen species (ROS) detection studies

#### Dihydrorhodamine 123 (DHR) and Dihydroethidium staining (DHE) staining

Tracheal cryosections (5μm) were incubated with 10 μM DHR or 5 μM DHE for 20 minutes at room temperature (RT). Cryosections Invitrogen™ ProLong™ Gold Antifade Mountant with DAPI (Invitrogen #P36941). Images were taken immediately, with identical exposure time for all settings using a confocal microscope (Leica TCS SP8 or Zeiss 800).

#### ROS detection assay

The ROS Detection Assay Kit (Abcam #ab287839) was used as per the manufacturer’s protocol. In brief, mouse lung mφs were treated with increasing iron-dextran concentrations. Cells were detached using detachment solution and centrifuged at 300 x g for 5 min at RT. Cells were stained with 1X ROS Label I/ROS Label for 30 minutes at 37°C and analyzed by flow cytometer (BD® LSR II).

### Lysosomal loss and acidification studies

#### Lysotracker Labeling

Mouse lung mφs were treated with Lysotracker^TM^ Green DND-26 (Invitrogen^TM^ #L7526) according to the manufacturer’s instructions. Cells were then analyzed by flow cytometry (BD LSR II, Agilent NovoCyte Quanteon, Agilent NovoCyte Penteon, BD^®^ FACSymphony^TM^) for median fluorescence intensity (MFI) levels by first gating on all cell material except small debris in the origin of a FSC versus SSC dot plot. H_2_O_2_ treated cells were used as a positive control.

#### Lysosomal acidification detection

Mouse lung mφs were placed in sterile coverslips and allowed to adhere by incubating at 37° C with 5% CO2 overnight. To detect lysosomal acidification LysoSensor^TM^ Green DND-189 probe (Invitrogen^TM^ #L7535) was used according to manufacturer’s instructions. Iron dextran and LysoSensor^TM^ were added to each sample. Cells were then incubated for 1h at 37° C with 5% CO2. Cells were washed, and images were taken using a confocal microscope (Leica TCS SP8, Leica STELLARIS).

#### Cellular pH measurement

To measure the pH of cellular organelles, we used LysoSensor™ Yellow/Blue dextran (Invitrogen^TM^ #L22460). To form a standard curve, lung mφs were treated with increasing pH media (1 to 8) for 1 h at 37°C. Lung mφs were treated with increasing concentrations of iron-dextran (0.04 mg/mL-1.6 Cells were then washed with PBS. The cell fluorescence was captured by imaging (Leica TCS SP8, Leica STELLARIS) and plate reader (Agilent BioTek Synergy multimode reader). Images were taken using a confocal microscope at excitation 405nm and 450nm and emission 505 and 530nm respectively. Plate reader readings included repetitive sweeps of the plate and alternating excitation as above. Fluorescence intensity from images was quantified by Image J. pH was calculated based on the standard curve.

#### Lysosomal staining

Mouse lung mφs were added in sterile coverslips and allowed to adhere by incubating at 37° C with 5% CO2 overnight. Iron dextran treatments were added for 16h. Cells were fixed with 4% PFA and permeabilized with 0.1% Triton X-100. Cells were incubated with 1% BSA, 22.52 mg/mL glycine in PBST for 30 min to block nonspecific staining sites. Lysosomes were stained with anti-LAMP1 Ab (Invitrogen # NBP225183P) for 1h at RT. Cells were washed with PBS and mounted using Invitrogen™ ProLong™ Gold Antifade Mountant with DAPI. Images and z stacks were taken using a confocal microscope (Leica TCS SP8 and Leica STELLARIS).

#### Phagolysosomal leakage assay

Cells were plated at appropriate density treated with 1mg/ml 4-kDa FITC-dextran (Sigma-Aldrich #46944) or 4-kDa TRITC-dextran (Sigma-Aldrich #T1037) in complete media for 2.5 hours at 37°C. Cells were washed with PBS and increasing iron dextran concentrations (0.04 mg/ml-1.6 mg/ml) were added. Cells were fixed with 4% PFA at 15, 30 and 60 min. Cells were mounted using Invitrogen™ ProLong™ Gold Antifade Mountant with DAPI. Images and z stacks were taken using a confocal microscope (Leica TCS SP8 and Leica STELLARIS).

### Cellular assays

#### Cell viability assays

Cell viability was assessed by the qualitative sulforhodamine B (SRB) assay kit (CYTOSCAN™ SRB G-Biosciences #786213) as per manufacturer’s instructions. Absorbance was measured at 565nm with microplate reader (Agilent BioTek Synergy multimode reader).

#### qPCR with reverse transcription

Total RNA was extracted using the RNeasy Plus Mini Kit (50) (Qiagen #74134) as per manufacturer’s instructions. 1μg RNA was reverse transcribed with Superscript II cDNA synthesis kit (Invitrogen, #18064014). Quantitative PCR (qPCR) was conducted in triplicate in a 20 μL reaction mixture using the PowerUPTM SYBRTM Green Master Mix for qPCR (Applied BiosystemsTM #A25742). in a CFX Opus Real-Time PCR System (BIO-RAD). Data were analyzed by the 2–ΔΔCt method, normalized to GADPH, and compared with controls.

#### Human studies

Samples from a total of 26 patients (12 males and 14 females; aged 22–70 years; average age, 57.5±15 years) with lung transplant (n=15) or without (control, n=11) were collected and analyzed. BALs were obtained from a convenience sample of patients undergoing bronchoscopy as part of their usual clinical care. Only adults >18 years old were included in the study. Where nodules were present as the reason for bronchoscopy, the BAL was performed on the contralateral side. Lavage was performed prior to any planned biopsy procedure. Patients with suspected fungal infection, as the primary reason for the bronchoscopy were excluded from the study.

#### Isolation of human alveolar mφs (AMs) from BAL

BAL fluid (5 - 20 mL) was centrifuged at 250 x g at 4 °C for 10 minutes. The cell pellet was washed with 20 mL wash buffer (PBS + 2% Heat inactivated FBS + 2 mM EDTA) and centrifuged at 250 x g at 4 °C for 10 minutes. BAL cells were resuspended in 100 μL of sorting buffer (PBS + 2% Heat inactivated FBS + 1 mM EDTA + 25 mM HEPES). Nonspecific antibody staining was blocked using human Fc block (BD Biosciences #564220) for 20 minutes on ice. AMs were stained for markers (anti-human CD45 BioLegend #304017, CD11b BioLegend #101216, HLA-DR BioLegend #307618, CD169 BioLegend #346008, CD206 BioLegend #321106, CD163 BioLegend #333612). Human AMs were sorted (BD FACSAria^TM^ II cell sorter) directly into 1 mL human AM culture medium (RPMI 1640 media supplemented with 10% FBS, 20% L-929 culture supernatant (with mφ colony stimulating factor, 100 U/mL final concentration), 1 mM sodium pyruvate, 10 mM N-2-hydroxyethylpiperazine-N’-2-ethanesulphonic acid (HEPES), and 1x penicillin/streptomycin.

#### In vitro culture of human AMs

Trypan blue was used to identify dead cells during cell counting. Viable AMs were plated at 1 x 10^5^/mL in 6 well plates in human AM culture media (see above). Cells were incubated in a 37°C humidified incubator with 5% CO2 atmosphere.

#### Phospholipid assay

To detect the phospholipid concentration in human BAL samples we used the Phospholipid Assay Kit (Sigma-Aldrich #MAK122), as per manufacturer’s protocol. Absorbance was measured at 570nm using a microplate reader (Agilent BioTek Synergy multimode reader). The concentration of phospholipids in the sample was calculated using the equation: µM = [(M_sample_ – M_blank_)/Slope], M_sample_ = Absorbance or fluorescent intensity measured in unknown sample, M_blank_ = Absorbance or fluorescent intensity measured in blank, Slope = Determined from standard curve (µM^-1^)

#### Lipid peroxidation assay

To measure the hydroperoxides directly from redox reactions with ferrous ions in the BAL supernatant we used the lipid peroxide (LPO) assay kit (Cayman Chemical #705002), as per manufacturer’s protocol. Absorbance was measured at 500nm using a microplate reader (Agilent BioTek Synergy multimode reader). The concentration of hydroperoxide was calculated using the equation: Hydroperoxide concentration (µM) = (HPST/VE) x (1ml/SV), VE (ml) = sample volume used for the assay, SV (ml) = original sample volume

#### 4-Hydroxynonenal (4-HNE) assay

4-HNE was measured using the 4-HNE ELISA Kit (NOVUS biologicals #NBP2-66364), as per manufacturer’s protocol. Absorbance was measured at 570nm using a microplate reader (Agilent BioTek Synergy multimode reader). The concentration of 4-HNE in the samples was calculated from the standard curve.

#### Malondialdehyde (MDA) assay

MDA was measured using the Malondialdehyde ELISA Kit (NOVUS biologicals #NBP2-78753), as per manufacturer’s protocol. Absorbance was measured at 570nm using a microplate reader (Agilent BioTek Synergy multimode reader). The concentration of MDA in the samples was calculated from the standard curve.

#### Iron assay from BAL

Total BAL iron was measured using the iron assay kit (Abcam #ab83366), as per manufacturer’s protocol. Absorbance was measured at 593nm using a microplate reader (Agilent BioTek Synergy multimode reader). Total iron concentration was calculated using the equation: Iron concentration = (S_a_/S_v_), S_a_ = content of iron in sample well calculated from standard curve. S_v_ = volume of sample added into the reaction wells

### Statistical analysis

All data shown are the mean±SEM, or mean±SD, and the number (n) in each figure legend represents biological or technical replicates, as specified. All experiments (except those described otherwise in the legend) were performed independently at least twice. For mouse experiments, at least five animals were included per group. Two-tailed Student’s *t* tests and one-way or two-way ANOVA followed by Bonferroni’s, Dunnett’s, Tukey’s or Sidak’s multiple-comparison tests were performed using GraphPad Prism 10 (GraphPad Software) (see figure legends for more details). The results of the statistical analyses are presented in each figure. *P*<0.05 was considered to be statistically significant. Association statistics of ferroptosis markers with conidial killing and fungal and bacterial infections were analyzed with univariate regression models and multivariate regression models. Covariates included all ferroptotic markers, age, and daily dosing of tacrolimus, prednisone, and MMF. P-values ≤ 0.05 for beta coefficients in the regression models were considered significant.

#### Transcriptome studies

RNA was extracted, and library preparation was performed, using a previously published BAL pipeline.(35) Briefly, 200 uL of unprocessed BAL was thawed, combined with 200 uL bead-bashing at 30Hz (TissueLyser II, Qiagen) with 60 seconds rest on ice between each cycle. Samples were centrifuged for 10 min at 4 and supernatant used for magnetic bead-based RNA extraction (Zymo Quick-RNA kit). Result RNA underwent cDNA library preparation (New England Biolabs Ultra II RNA Library Prep Kit), and PCR-amplified libraries sequenced to a target depth of 20,000 read-pairs per sample on a NovaSeq S4 instrument (Illumina). Resultant fastq files were filtered for quality and complexity and aligned to human genome v38 to generate transcript counts (STAR). Non protein-coding, mitochondrial, and ribosomal reads were excluded and resulting protein-coding transcript counts underwent gene set variance analysis (GSVA), using Reactome and Gene Ontology: Biological Function repositories to produce semi-normally distributed gene set enrichment scores. Gene set enrichment scores were compared between lung transplant recipients vs. controls using the Wilcoxon rank-sum test and correlated with BAL measures of conidial killing and markers of ferroptosis using the non-parametric Spearman correlation. Data from the “Gene Expression Profiling of Bronchoalveolar Lavage Cells During *Aspergillus* Colonization of the Lung Allograft” study were accessed via ebi.ac.uk (E-MTAB-6040).(36) This dataset includes adult lung transplant recipients colonized with *Aspergillus*. GSVA was applied to generate gene set enrichment scores to the Reactome collection. Genes differentially expressed (fold change ≥ 2 and P < 0.05) in *Aspergillus* colonization cases compared to infection-free controls (36) were subjected to functional enrichment analysis. Upregulated and downregulated gene sets were analyzed in gene ontology (GO) consortium resources (https://geneontology.org/;release: 2025-02-06:40, 267)(53, 54), using the PANTHER analysis tool.(55, 56) Significant (P < 0.05) GO aspects (biological process) and reactome pathways (Reactome version 86 Released 2023-09-07) associated with iron metabolism, ROS and ferroptosis were retrieved and their background frequency, sample frequency, indication of over/underrepresentation were studied.

#### Reagents

RAS-selective lethal (RSL3, Cayman #19288), ferrostatin-1 (Sigma #SML0583), chlodronate liposomes (Encapsula Nanaoscience #CLD-8909), BOPDIPY^TM^ 581/591 C11 (Invitrogen™ #D3861), iron dextran (Sigma (Fisher Bioreagents # BP531-25), Triton X-100 (Thermo Scientific Chemicals #A16046-AE), DMSO (Corning #25-950-CQC), Tween 80 (Millipore Sigma #655207), Tween 20 (Millipore Sigma #655206).

## Author Contributions

EIM, AE-H, HS, PC, MZ, PJ, AE, and JLH designed research studies, conducted experiments, acquired and analyzed data, and were involved in writing and editing the manuscript. ED, WC, JQ, BS, JC, SP, CHO, and SO acquired and analyzed data and were involved in editing the manuscript. GD acquired and analyzed data, helped design experiments and was involved in editing the manuscript. BG, JF, JMV were involved in the study design and conduct of experiments, provided reagents, and in writing and editing the manuscript.

## Acknowledgements

*Funding.* JLH is funded by R01HL157414-01-NIH/NHLBI. MZ is funded by K23HL146936-NIH/NHLBI. JMV is funded by R01AI150181-NIH/NIAID. BG is funded by R35GM137936-NIH. JF is funded by R01AI158442-NIH/NIAID, R01AI143197-NIH/NIAID, and R21AI178048-NIH/NIAID. Data/sorting were collected and performed on instruments in the Shared FACS Facility obtained using NIH S10 Shared Instrument Grants S10RR027431-01, S10RR025518-01 and 1S10OD026831-01. We would like to thank all patients for their willingness to participate in the study. Cell sorting/flow cytometry analysis for this project was done on instruments in the Stanford Shared FACS Facility. We would like to acknowledge Dr. David Stevens for his support and for providing the *Af* isolates used in the study and Dr. Tobias Hohl for providing us with the *Af*-FLARE conidia. We would also like to thank Dr. Guang-Shing Cheng and Dr. Ajay Sheshadri for their careful review of the manuscript.

