## Supplemental Figure 1 for "Macrophage ferroptosis inhibits *Aspergillus* conidial killing in lung transplantation"

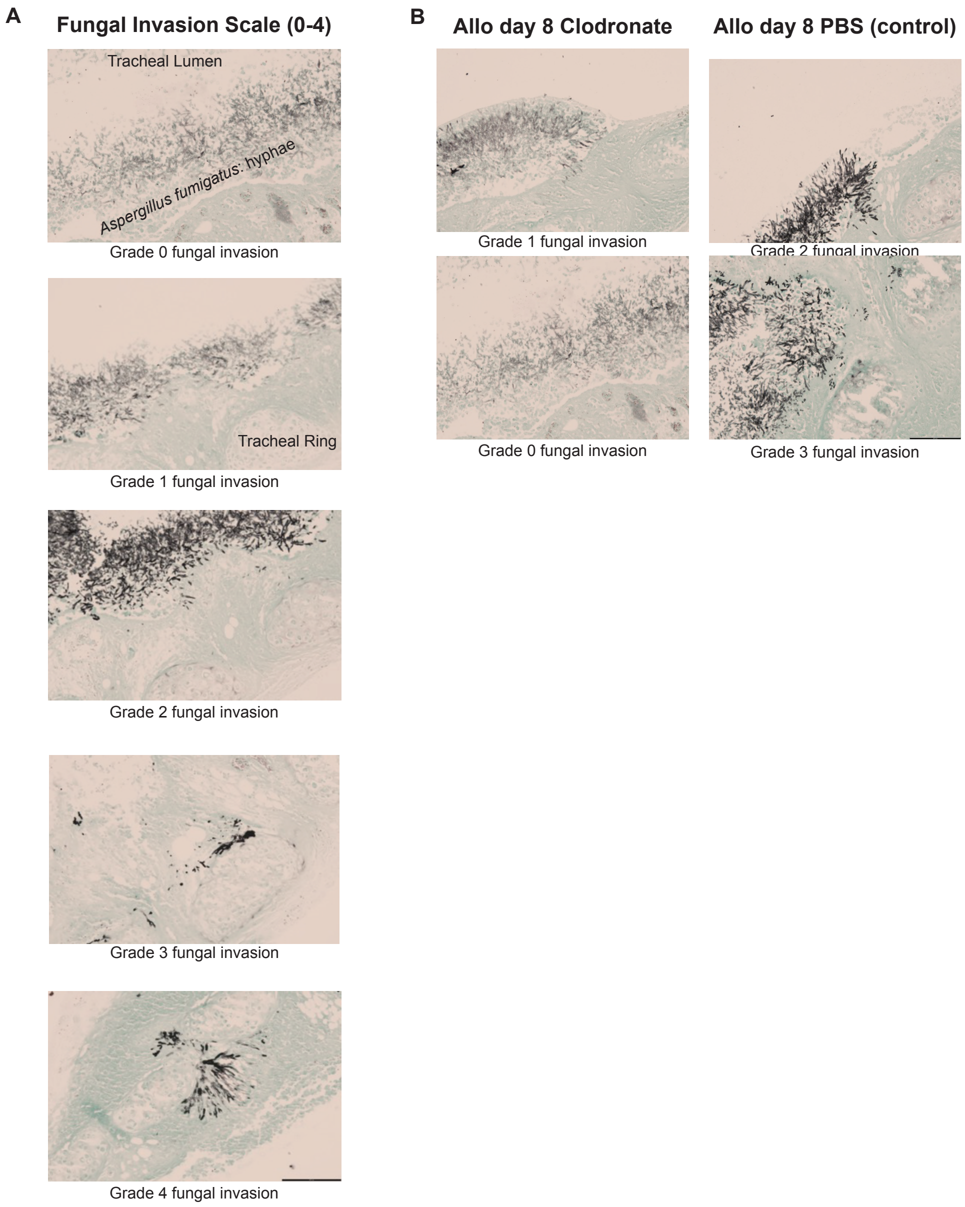


**Supplemental Figure 1. Fungal Invasion in orthotopic tracheal transplant model of *Aspergillus fumigatus* infection** (**A**) Grocott's Methenamine Silver (GMS) staining of explanted orthotopic tracheal transplant (OTT) depicting semiquantitative scale of fungal invasion: Grade 0, no invasion; Grade 1: invasion of epithelial layer; Grade 2: invasion of sub-epithelial layer; Grade 3 invasion to the depth of the tracheal ring' and Grade 4 invasion beyond the tracheal ring. (12-14) (**B**) Representative images of explanted GMS stained trachea for allotransplants day 8, clodronate liposome (macrophage depleted) treated animals (right panels) and PBS liposome (control) treated animals (left panels).
