## Supplemental Figure 2 for "Macrophage ferroptosis inhibits *Aspergillus* conidial killing in lung transplantation"

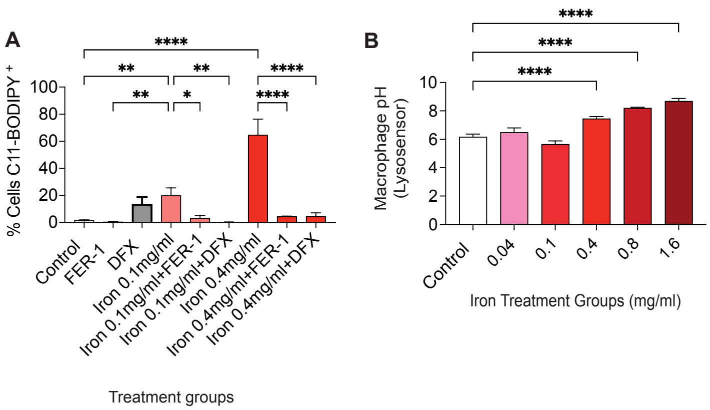


**Supplemental Figure 2. In vitro ferroptosis and macrophage pH studies** (**A**) Percent cells C11-BODIPY positive (marker of ferroptosis) by treatment group. Primary murine macrophages co-cultured with iron (0.1 and 0.4mg/ml) alone or with ferrostatin-1 (FER-1, ferroptosis inhibitor) or deferasirox (DFX, iron chelator). Data as analyzed by one way ANOVA with Tukey's correction for multiple comparisons. Data represent mean +/- SD. (n = 3 biologic and technical replicates/group). (**B**) Macrophage pH as measured by ratio of LysoSensor yellow to blue dextran fluorescent probes. Data as analyzed by one way ANOVA with Dunnett's correction for multiple comparisons. (n = 3 biologic and technical replicates/group). Data presented are mean +/- SEM. *$P < 0.05$, **$P < 0.01$, ***$P < 0.001$, ****$P < 0.0001$.
