## Supplemental Figure 3 for "Macrophage ferroptosis inhibits *Aspergillus* conidial killing in lung transplantation"

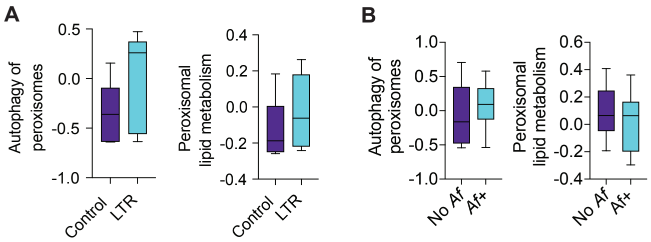


**Supplemental Figure 3. In vitro ferroptosis and macrophage pH studies** (**A**) The expression of the BAL Reactome gene sets "Autophagy of peroxisomes" and "Peroxisomal lipid metabolism" comparing lung transplant recipients to non-lung transplant recipients **(B)** Gene set variance analysis of LTR colonized with *Af* (*Af*+, n=12) and LTR without evidence of *Af* colonization (No *Af*, n=10) accessed from ebi.ac.uk (36), depicting "Autophagy of peroxisomes" and "Peroxisomal lipid metabolism." Analyzed using the Mann-Whitney U test. Data presented are median and high and low.
