## Supplemental Tables for "Macrophage ferroptosis inhibits *Aspergillus* conidial killing in lung transplantation"

### Supplementary Material - Tables

#### Supplementary Table 1. Clinical Characteristics of Study Participants

| Patient Number | Sex | Age | Race/ ethnicity | Transplant Status and Indication | Time from bronchoscopy to transplant/ primary diagnosis (months) | Immune suppression at time of bronchoscopy | Antifungal prophylaxis/ treatment | Iron supplementation | Concomitant infections on bronchoalveolar lavage | Additional Clinical Details |
|---|---|---|---|---|---|---|---|---|---|---|
| 1 | F | 50-60 | Black | Heart, NICM | < 1 (heart transplant) | Tacrolimus, Prednisone, MMF | Caspofungin, Itraconazole | None | Varicella zoster virus | Extracorporeal membrane oxygenation (ECMO) for disseminated varicella |
| 2 | F | 40-50 | White (non-Hispanic) | Heart/Lung, Sjogren's disease, Pulmonary arterial hypertension (PAH) | 4 | Tacrolimus, Prednisone, MMF | Isavuconazole, Caspofungin | None | Vancomycin-resistant Enterococcus faecium | History of invasive mold infection of tracheal anastomosis |
| 3 | F | 60-70 | Asian | Lung, interstitial lung disease (ILD) Rheumatoid arthritis | 6 | Tacrolimus | Isavuconazole, | None | COVID-19 | COVID pneumonia on ECMO |
| 4 | M | 70-80 | White (non-Hispanic) | None | < 1 (s/p CABG for NSTEMI | Prednisone | Caspofungin | None | Staphylococcus aureus | Staphylococcus aureus pneumonia |
| 5 | F | 50-60 | Hispanic | Lung, ILD | 29 | Tacrolimus, Prednisone | Isavuconazole | None | COVID pneumonia, Burkholderia cepacia (colonization) | COVID pneumonia and ARDS on ECMO after transplant |
| 6 | M | 60-70 | Hispanic | Lung, Hypersensitivity pneumonitis | < 1 | Tacrolimus, Prednisone | Isavuconazole | None | None | |
| 7 | M | 60-70 | White (non-Hispanic) | Lung, ILD | 6 | Tacrolimus, Prednisone | Posaconazole | None | Klebseilla oxytoca | |
| 8 | F | 30-40 | Black | Lung, ILD dermatomyositis | 12 | Tacrolimus, Prednisone | Posaconazole | Yes | Stenotrophomonas maltophilia, Serratia rubidaea | |
| 9 | M | 40-50 | Asian | Lung/ Kidney, ILD, scleroderma | 9 | Tacrolimus, Prednisone | Isavuconazole | None | Pseudomonas aeruginosa | |
| 10 | M | 60-70 | White (non-Hispanic) | Lung, ILD | 15 | Tacrolimus, Prednisone | Posaconazole | None | Staphylococcus aureus, Steprococcus agalactiae | |
| 11 | M | 60-70 | Hispanic | Lung, Hypersensitivity pneumonitis | 15 | Tacrolimus, Prednisone, Everolimus | Posaconazole | Yes | Klebseilla oxytoca, Stenotrophophomas maltophilia | |
| 12 | M | 50-60 | White (non-Hispanic) | Lung, COVID-19 | 18 | Tacrolimus, Prednisone | Posaconazole | Yes | COVID-19 | |
| 13 | M | 40-50 | Asian | Lung, ILD dermatomyositis | 18 | Tacrolimus, Prednisone, Everolimus | Isavuconazole, Caspofungin | Yes | Pseudomonas aeruginosa | |
| 14 | F | 60-70 | White (non-Hispanic) | Heart, NICM | < 1 | Tacrolimus, Prednisone, MMF | Posaconazole, Caspofungin | None | Staphylococcus aureus | ECMO for ARDS |
| 15 | F | 60-70 | Other | None, cardiac arrest | < 1 | None | None | None | None | ECMO for cardiogenic shock |
| 16 | M | 60-70 | Asian | None, s/p AVR, CABG | 1 | None | None | Yes | Stenotrophas maltophilia | ECMO for ARDS |
| 17 | F | 40-50 | Asian | None, cardiac arrest | 1 | None | None | None | Escherichia coli | ECMO after cardiac arrest |
| 18 | M | 50-60 | Black | Lung, ILD, Sarcoidosis | 12 | Tacrolimus, Prednisone, Everolimus | Posaconazole | Yes | None | |
| 19 | F | 20-30 | Hispanic | Lung, Cystic fibrosis, retransplant | 6 | Tacrolimus, Prednisone, MMF | Isavuconazole | None | None | Cystic fibrosis retransplanted |
| 20 | F | 60-70 | Hispanic | Lung, ILD, Sjogren's | < 1 | Tacrolimus, Prednisone, MMF | None | None | None | Transplant complicated by ischemia reperfusion injury |

| | | | | | | | | | | |
|---|---|---|---|---|---|---|---|---|---|---|
| 21 | M | 60-70 | White (non-Hispanic) | Lung, PAH | 25 | Tacrolimus, Prednisone | Posaconazole | None | None | |
| 22 | F | 60-70 | White (non-Hispanic) | None, vaginal cancer | 18 (vaginal cancer) | None | None | None | None | History of metastatic vaginal cancer |
| 23 | F | 60-7- | Asian | None, Benign nasopharyngeal mass | 14 (nasopharyngeal mass) | None | None | None | None | |
| 24 | F | 50-60 | White (non-Hispanic) | Heart | 189 (heart transplant) | Tacrolimus, Sirolimus | None | None | None | Heart transplant, metastatic cervical cancer undergoing treatment |
| 25 | F | 20-30 | Asian | None, Cryptogenic organizing pneumonia (COP) | <1 (COP) | None | None | None | None | Cryptogenic organizing pneumonia |
| 26 | F | 60-70 | White (non-Hispanic) | None, metastatic ovarian cancer | 41 (ovarian cancer) | None | None | None | None | Metastatic ovarian cancer |

**Abbreviations: AVR**: Aortic valve replacement, **CABG**: Coronary artery bypass graft, **ECMO**: Extracorporeal Membrane Oxygenation, **ILD**: Interstitial lung disease, **MMF**: Mycophenolate mofetil, **NICM:** Non-ischemic cardiomyopathy, **PAH**: Pulmonary arterial hypertension.

**Supplementary Table 2. Univariate regression analysis for conidial killing, fungal infection, and bacterial infection**

| | Conidial killing (0 – 100%) | Fungal infection (0 – 4 scale) | Bacterial infection (0 – 4 scale) |
|---|---|---|---|
| **MDA (beta, p-value)** | -0.270, 0.00001 | 0.918, 0.00923 | 0.941, 0.00345 |
| **4-HNE (beta, p-value)** | -0.141, 0.00001 | 0.469, 0.00231 | 0.483, 0.000505 |
| **LPO (beta, p-value)** | -0.002, 0.00025 | 0.009, 0.00751 | 0.008, 0.00492 |
| **Iron (beta, p-value)** | -2.668, 0.0074 | 5.084, 0.324 | 7.123, 0.131 |

**4-HNE: 4-Hydroxynonenal, LPO: lipid hydroperoxide; MDA: malondialdehyde**

**Supplementary Table 3: Multivariate regression for conidial killing for MDA, 4-HNE, LPO, and iron adjusted for daily dose of immune suppressive medications and age**

| | MDA | 4-HNE | LPO | Iron |
|---|---|---|---|---|
| **Other markers of ferroptosis (beta, p-value)** | -0.261, 0.00137 | -0.136, < 0.00001 | -0.002, 0.00450 | -2.483, 0.0343 |
| **Tacrolimus daily dosing (beta, p-value)** | -0.020, 0.43315 | -0.036, 0.0171 | -0.040, 0.13089 | -0.058, 0.0501 |
| **Prednisone daily dosing (beta, p-value)** | -0.002, 0.77396 | $-2.280 \times 10^{-4}$, 0.9429 | $1.688 \times 10^{-3}$, 0.78968 | -0.0004, 0.9507 |
| **Mycophenolate Mofetil daily dosing (beta, p-value)** | $3.931 \times 10^{-6}$, 0.97137 | $3.442 \times 10^{-5}$, 0.5783 | $1.472 \times 10^{-5}$, 0.90053 | 0.0001, 0.2994 |
| **Age (beta, p-value)** | $2.567 \times 10^{-3}$, 0.53837 | $1.145 \times 10^{-3}$, 0.6154 | $3.386 \times 10^{-4}$, 0.93633 | -0.002, 0.5842 |

**Supplementary Table 4: Multivariate regression for fungal infection for MDA, 4-HNE, LPO, and iron each adjusted for daily dose of immune suppressive medications and age**

| | MDA | 4-HNE | LPO | Iron |
|---|---|---|---|---|
| **Other markers of ferroptosis (beta, p-value)** | 0.853, 0.0626 | 0.452, 0.00953 | 0.009, 0.0309 | 5.163, 0.4151 |
| **Tacrolimus daily dosing (beta, p-value)** | 0.053, 0.7432 | 0.104, 0.44104 | 0.100, 0.4922 | 0.181, 0.2644 |
| **Prednisone daily dosing (beta, p-value)** | 0.007, 0.8323 | 0.003, 0.93523 | -0.011, 0.7551 | 0.013, 0.7521 |

| | | | | |
|---|---|---|---|---|
| **Mycophenolate Mofetil daily dosing (beta, p-value)** | -0.0008, 0.2650 | -0.0009, 0.15297 | -0.0007 0.2974 | -0.001, 0.0952 |
| **Age (beta, p-value)** | -0.006, 0.8060 | -0.002, 0.92385 | -0.003, 0.8932 | 0.0129, 0.6224 |

**Supplementary Table 5: Multivariate regression for bacterial infection for MDA, 4-HNE, LPO, and iron each adjusted for daily dose of immune suppressive medications and age.**

| | **MDA** | **4-HNE** | **LPO** | **Iron** |
|---|---|---|---|---|
| **Other markers of ferroptosis (beta, p-value)** | 1.101, 0.0117 | 0.524, 0.0015 | 0.010, 0.00774 | 9.588, 0.118 |
| **Tacrolimus daily dosing (beta, p-value)** | $-5.470 \times 10^{-3}$, 0.9701 | 0.072, 0.5499 | 0.065, 0.62569 | 0.155, 0.309 |
| **Prednisone daily dosing (beta, p-value)** | -0.020, 0.5185 | -0.023, 0.4071 | -0.040, 0.23710 | -0.022, 0.564 |
| **Mycophenolate Mofetil daily dosing (beta, p-value)** | $-1.277 \times 10^{-5}$, 0.9838 | -0.0002, 0.7263 | $2.707 \times 10^{-5}$, 0.96491 | -0.0006, 0.409 |
| **Age (beta, p-value)** | -0.016, 0.5050 | -0.008, 0.6920 | $-9.601 \times 10^{-3}$, 0.66567 | 0.006, 0.799 |

**Supplementary Table 6: Gene ontology biological processes and reactome pathways associated with ferroptosis and over-represented in LTR with *Aspergillus* colonization.** (36)

| **Enrichment terms** | **Background frequency (genes from the human genome)** | **Sample frequency (genes upregulated in the disease gene set)** | **Sample gene names** | **Fold change** | **Unadjusted P-value** | **False discovery rate (FDR)** |
|---|---|---|---|---|---|---|
| **Iron ion transport (GO:0006826)** | 54 | 7 | SLC40A1, SLC39A8, STEAP4, SLC25A37, LTF, LCN2, CP | 6.84 | 0.0000684 | 0.00638 |
| **Transition metal ion transport (GO:0000041)** | 101 | 9 | TCN1, TMEM163, SLC40A1, SLC39A8, STEAP4, SLC25A37, LTF, LCN2, CP | 4.7 | 0.000129 | 0.0108 |
| **Programmed cell death (GO:0012501)** | 1094 | 42 | YAP1, PFKFB3, IL1B, GULP1, CD24, RASSF6, TNFRSF10C, TOX3, NR4A2, SNCA, PRKCB, MECOM, SLC40A1, NLRP3, KRT18, MARCKS, TNFSF10, CAV1, ERBB3, THRB, BNIP3, ZFP36L1, WNT5A, DAPL1, PLS3, STK26, IGFBP3, PLS1, CXCR4, KITLG | 2.03 | 0.0000171 | 0.00233 |
| **Cell death (GO:0008219** | 1099 | 42 | YAP1, PFKFB3, IL1B, GULP1, CD24, RASSF6, TNFRSF10C, TOX3, | 2.02 | 0.000018 | 0.00243 |

| | | | | | | |
|---|---|---|---|---|---|---|
| | | | NR4A2, SNCA, PRKCB, MECOM, SLC40A1, NLRP3, KRT18, MARCKS, TNFSF10, CAV1, ERBB3, THRB, BNIP3, ZFP36L1, WNT5A, DAPL1, PLS3, STK26, IGFBP3, PLS1, CXCR4, KITLG | | | |
| **Response to lipid (GO:0033993)** | 856 | 33 | DGAT2, YAP1, FFAR2, IL1B, ALPL, SLPI, PDE4B, SPP1, DSG2, SNCA, PRKCB, NLRP3, ANXA3, GNG2, CAV1, CXCL6, AREG, FCAR, AKR1C1, ZFP36L1, WNT5A, PLS3, CXCL8, PELI1, ADM, YES1, FOXA1, FGFR2, IGFBP7, CX3CR1 | 2.03 | 0.000151 | 0.0118 |
| **Response to oxygen-containing compound (GO:1901700)** | 1585 | 50 | DGAT2, YAP1, FFAR2, IL1B, PIK3R3, ALPL, SLPI, PDE4B, SPP1, DSG2, MMP12, NR4A2, ST6GAL1, SNCA, PRKCB, PTPRK, IGF1, NLRP3, TNFSF10, GNG2, JUP, CAV1, CXCL6, PTPRN2, AREG, DDR1, FCAR, AKR1C1, INAVA, MET | 1.66 | 0.000354 | 0.0211 |
